# oxo-flow: compiled, memory-safe bioinformatics workflow orchestration

**DOI:** 10.64898/2026.06.11.731578

**Authors:** Shixiang Wang

## Abstract

Bioinformatics analyses depend on workflow engines to coordinate dozens of computational tools across complex dependency chains. The most widely adopted engines—Snakemake, Nextflow, the Common Workflow Language (CWL), and the Workflow Description Language (WDL)—run on interpreted or just-in-time (JIT) compiled language runtimes, incurring hundreds of milliseconds of startup latency and providing no compile-time safety guarantees from the host language. We developed oxo-flow, a workflow engine written in Rust that compiles to a single native binary. On an Apple M5 processor, oxo-flow parses, validates, and dry-runs a production-scale workflow in roughly 22 milliseconds—before Snakemake or Nextflow have finished loading their runtime environments. Peak memory usage is 16 megabytes, representing six- to seven-fold reductions relative to Snakemake and Nextflow. Dry-run latency is essentially independent of workflow size: a hundred-fold increase in rule count adds approximately 0.4 milliseconds. oxo-flow integrates 31 command-line tools, a REST interface with 60 endpoints, an embedded web application, and native cluster submission into a single 10-megabyte binary. It provides per-rule environment isolation across seven backends, checkpoint-based fault tolerance with cryptographic output verification, and a formal installation and operational qualification protocol for regulated laboratory environments. Ten curated workflows and three demonstration pipeline repositories are available. oxo-flow is freely available under Apache License 2.0 at https://github.com/Traitome/oxo-flow.

## 1 Introduction

A bioinformatics analysis from raw sequencing data to interpretable results passes through 20–100 computational steps [1, 2]. Each step invokes a specialised tool with its own software dependencies; a single pipeline may simultaneously demand incompatible Python versions and conflicting C libraries [3]. Orchestrating this correctly, reproducibly, and efficiently across laptops, institutional high-performance computing (HPC) clusters, and cloud instances is a genuine engineering challenge [4].

Workflow management systems address this by providing a declarative language for specifying computational steps and a runtime that executes them in dependency order within isolated environments. Several systems have achieved broad adoption. Snakemake [1,5] embeds a Python domain-specific language, with a large ecosystem including ATLAS for metagenome assembly [6] and grenepipe for variant calling [7]. Nextflow [2] provides a reactive dataflow model in Groovy on the Java Virtual Machine. The Common Workflow Language [8] and Workflow Description Language [9] are community standards that prioritise multi-engine portability, with documented applications in RNA-seq [10], genome assembly [11], and provenance tracking [12]. Galaxy [13] wraps workflow construction in a graphical interface [14]. Anderson *et al*. [15] provide a comparative evaluation framework. BLIT [16] addresses tool integration from within R.

These engines share a consequential design property: they execute on interpreted or JIT-compiled language runtimes. Startup latency runs from hundreds of milliseconds to seconds. Malformed configurations surface at runtime. No formal memory-safety guarantee is provided by Python, Groovy, or Java. Monitoring, reporting, and cluster management are delegated to separate tools, fragmenting the reproducibility chain.

Individual tools have demonstrated that a systems language brings concrete advantages to bioinformatics. GFFx [17] uses Rust compilation to accelerate genomic feature extraction by 10–80-fold. Phylo-rs [18] provides a memory-safe phylogenetic library. No existing workflow engine, however, has been built with compile-time safety guarantees.

oxo-flow fills this gap: a compiled, memory-safe engine delivering the complete workflow lifecycle—command-line interface (CLI), application programming interface (API), web interface, cluster submission, and reporting—as a single binary. Written in Rust (2024 edition) [19], it guarantees memory safety and data-race freedom at compile time. Workflows are declared in TOML (Tom’s Obvious, Minimal Language) [20] and validated against a JSON (JavaScript Object Notation) Schema. The engine constructs a directed acyclic graph (DAG) from rule declarations, computes a topological execution order, identifies parallel-ready rule groups, and dispatches them under resource constraints with deadlock detection. Its design was informed by input from 30 domain experts spanning bioinformatics, clinical oncology, HPC, security, and regulatory affairs [26]. We describe version 0.7.0 and report comprehensive benchmarks measured on identical hardware against Snakemake 9.22.0 and Nextflow 26.04.3.

## 2 Results

### 2.1 System overview

oxo-flow is organised as three crates sharing a common core library (Figure 1). The core (`oxo-flow-core`, 24,417 lines of Rust) contains the engine: DAG construction, configuration parsing with compile-time state validation, rule modelling (90 public fields), wildcard expansion, environment management, local and cluster execution, resource scheduling, checkpoint persistence, container packaging, and report generation. The CLI binary (`oxo-flow-cli`, 4,618 lines) exposes this through 31 subcommands. The web binary (`oxo-flow-web`, 6,554 lines) wraps the same core behind a REST API with an embedded interface. Release builds produce binaries of 10 MB (CLI) and 6.7 MB (web server). Cross-compilation targets x86_64, aarch64, and armv7 Linux with static musl linking.

**Figure 1.**
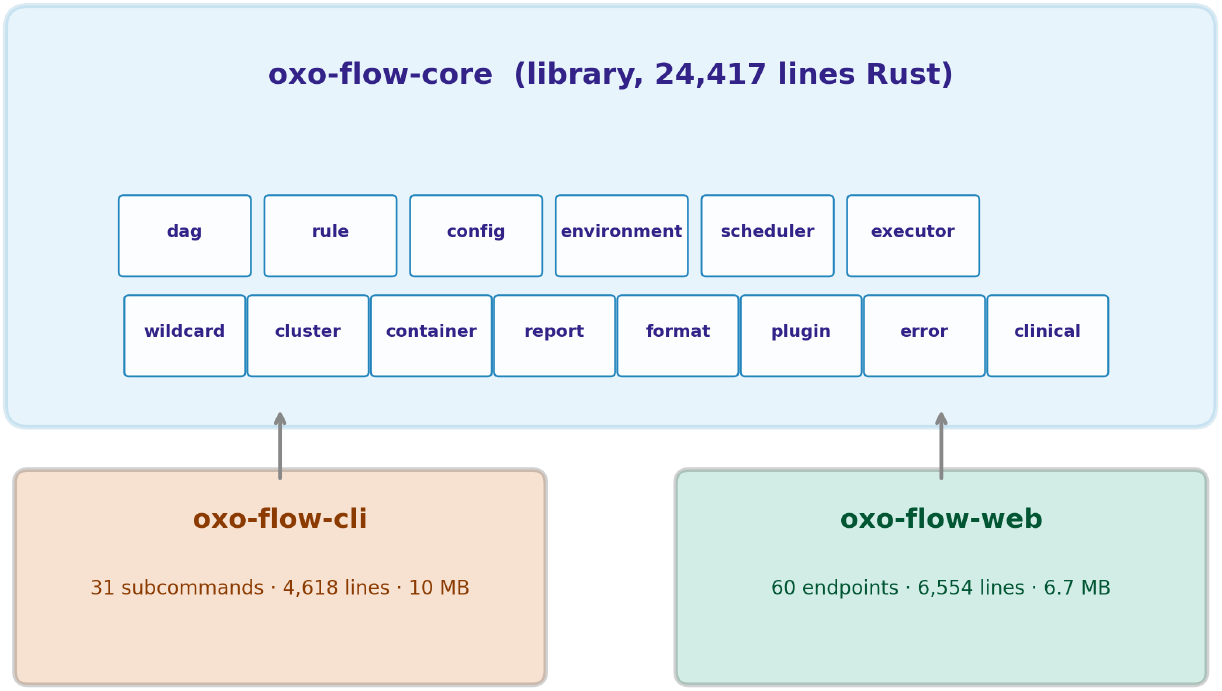
Three-crate workspace architecture. CLI and web binaries share the identical core library.

Workflows are specified in TOML and validated against a JSON Schema. The engine compiles each workflow into a directed acyclic graph with rules as nodes and data-flow dependencies as edges. From the DAG it derives topological execution order, parallel-ready rule groups (rules at identical depth with no intra-level dependencies), target-aware execution (the `-t` flag computes transitive dependency closure of specified outputs), and critical path analysis. Wildcard patterns with optional regex constraints support flexible sample-, read-, and chromosome-level iteration across independent dimensions.

Seven environment backends provide per-rule isolation: Conda, Pixi, Docker, Singularity, Venv, Modules, and the host system (Table 1). Each rule independently declares its environment; a resolver caches activations by content-addressed keys, eliminating redundant setup. Environment groups let related rules share a specification.

**Table 1:** Environment backends with cache-keyed activation.

| Backend | Specification | Cache key | Isolation |
| --- | --- | --- | --- |
| Conda | YAML environment file | YAML SHA-256 | Process |
| Pixi | `pixi.toml` | Lockfile hash | Project |
| Docker | Image name:tag | Image digest | Container |
| Singularity | SIF or definition file | Image hash | Container |
| Venv | `requirements.txt` | Requirements hash | Virtualenv |
| Modules | Module name/version | Module string | Shell env |
| System | Host environment | System fingerprint | None |

The executor dispatches rules as asynchronous subprocesses with semaphore-governed concurrency under user-specified parallelism limits and per-rule resource declarations. A resource pool tracks CPU, memory, GPU, and disk availability; deadlock detection distinguishes permanent stalls from temporary starvation. Checkpoint-based fault tolerance persists rule outcomes, benchmark records, configuration checksums, and output file hashes to JSON, enabling zero-redundancy recovery from interruptions.

A clinical reporting module generates structured reports with variant classification, biomarker tracking, and compliance audit trails, supporting regulated genomics environments. The web server exposes 60 REST endpoints with session-token authentication, per-IP rate limiting, and workspace-isolated state persistence. Cluster backends support SLURM, PBS, SGE, and LSF; container packaging generates self-contained Docker or Singularity images. A formal Installation Qualification / Operational Qualification / Performance Qualification (IQ/OQ/PQ) protocol template is provided for laboratories operating under regulatory frameworks.

### 2.2 Operation latency, scaling, and memory efficiency

We measured latency using hyperfine [25] on an Apple M5 MacBook Pro (10 cores, 24 GB unified memory, macOS 26.5.1). All measurements included warmup runs.

Binary startup completes in 6.9 ms (*σ* = 0.9 ms, *n* = 50). Full workflow dry-run—parsing, validating, constructing the DAG, and printing the execution plan—was benchmarked on all ten curated gallery workflows and on synthetic linear-chain workflows of 1, 5, 10, 20, 50, and 100 rules. Dry-run latency is essentially constant: mean 13.3 ms, range 13.0–13.7 ms (Figure 2b). A 100-fold increase in workflow size adds approximately 0.4 ms.

**Figure 2.**
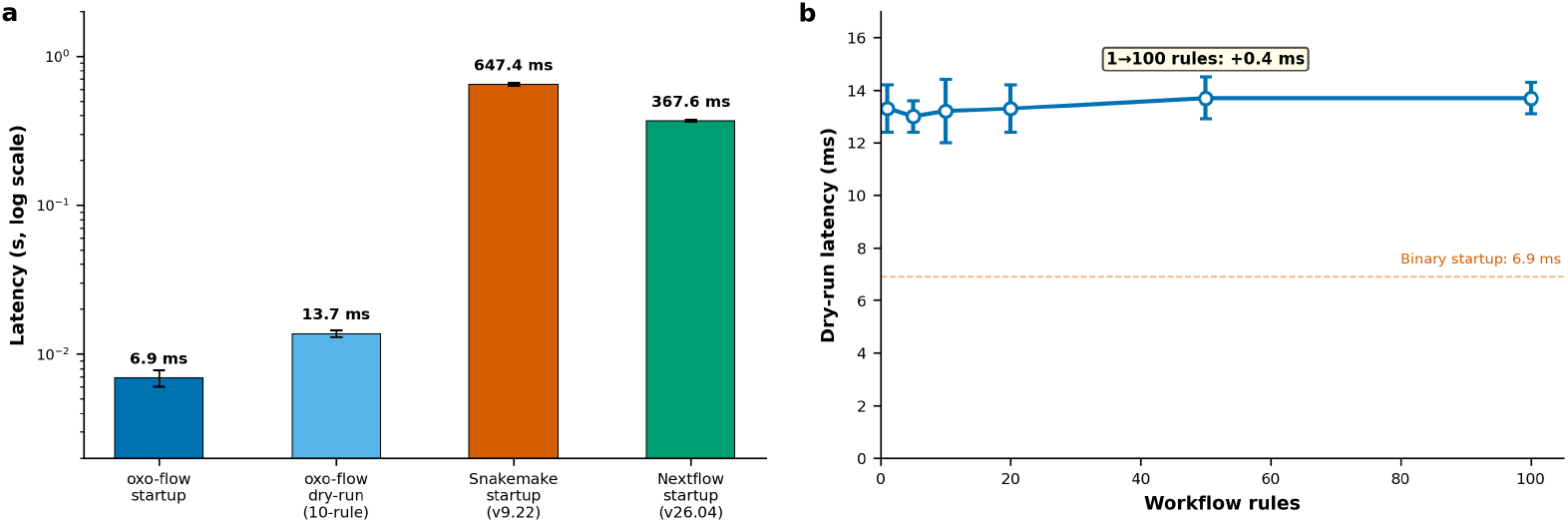
Operation latency and scaling. (a) Latency comparison across engines. (b) Dry-run latency on synthetic workflows of 1–100 rules.

For comparison, Snakemake 9.22.0 binary startup on identical hardware required 647.4 ms (*σ* = 15.0 ms, *n* = 20), and Nextflow 26.04.3 required 367.6 ms (*σ* = 4.7 ms, *n* = 15). oxo-flow’s complete dry-run cycle is completed before either engine has finished loading its runtime (Figure 2a). Matched dry-run benchmarks across DSLs were not performed; constructing semantically equivalent workflow definitions in different formats is non-trivial.

Peak resident set size was measured via `/usr/bin/time -l`. oxo-flow dry-run peaked at 16.4 MB. Snakemake startup peaked at 95.3 MB, and Nextflow at 121.0 MB (Figure 3c)—5.8*×* and 7.4*×* reductions. oxo-flow’s working set fits within the L3 cache of modern processors; the interpreted engines’ footprints are dominated by interpreter and JVM runtime overhead.

**Figure 3.**
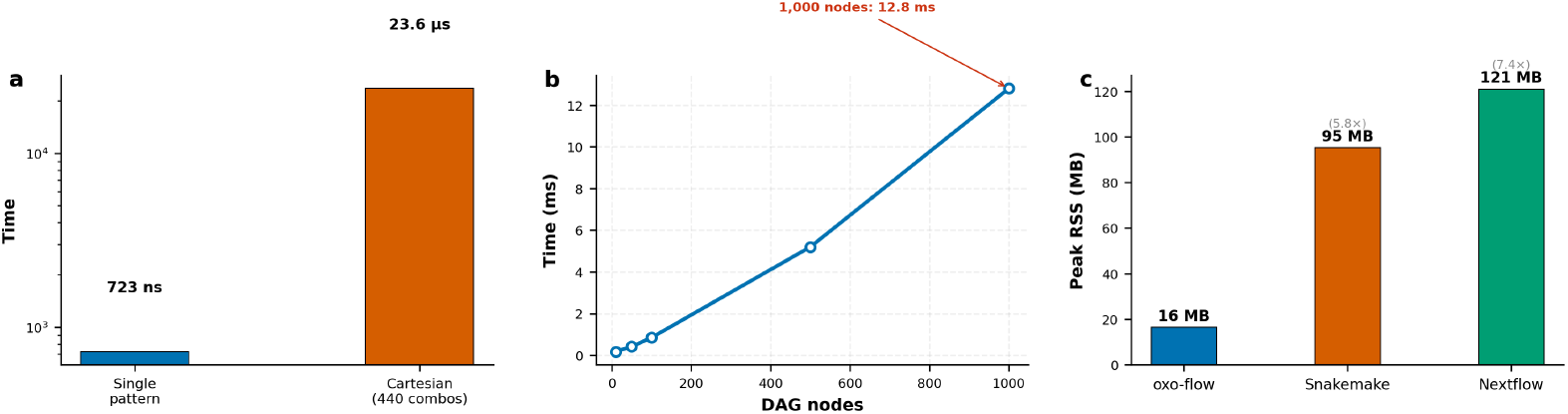
Component benchmarks and memory. (a) Wildcard engine. (b) DAG construction on linear chains. (c) Peak memory comparison.

### 2.3 Component benchmarks

Wildcard expansion and DAG construction were benchmarked independently (Figure 3a,b). Single-pattern expansion completes in 723 ns (Criterion.rs, *n* = 100). A Cartesian product of 440 combinations (10 samples *×* 2 reads *×* 22 chromosomes) completes in 23.6 *µ*s. DAG construction on linear chains scales approximately linearly with node count: 0.18 ms for 10 nodes, 0.85 ms for 100, and 12.8 ms for 1,000.

### 2.4 Workflow gallery and pipeline validation

oxo-flow provides 31 CLI subcommands spanning execution, quality assurance, project management, monitoring, environments, cluster, and utilities. Ten curated workflow examples are distributed with the software and validated in continuous integration, ranging from a single-rule hello-world to multi-omics integration (Table 2).

**Table 2:**
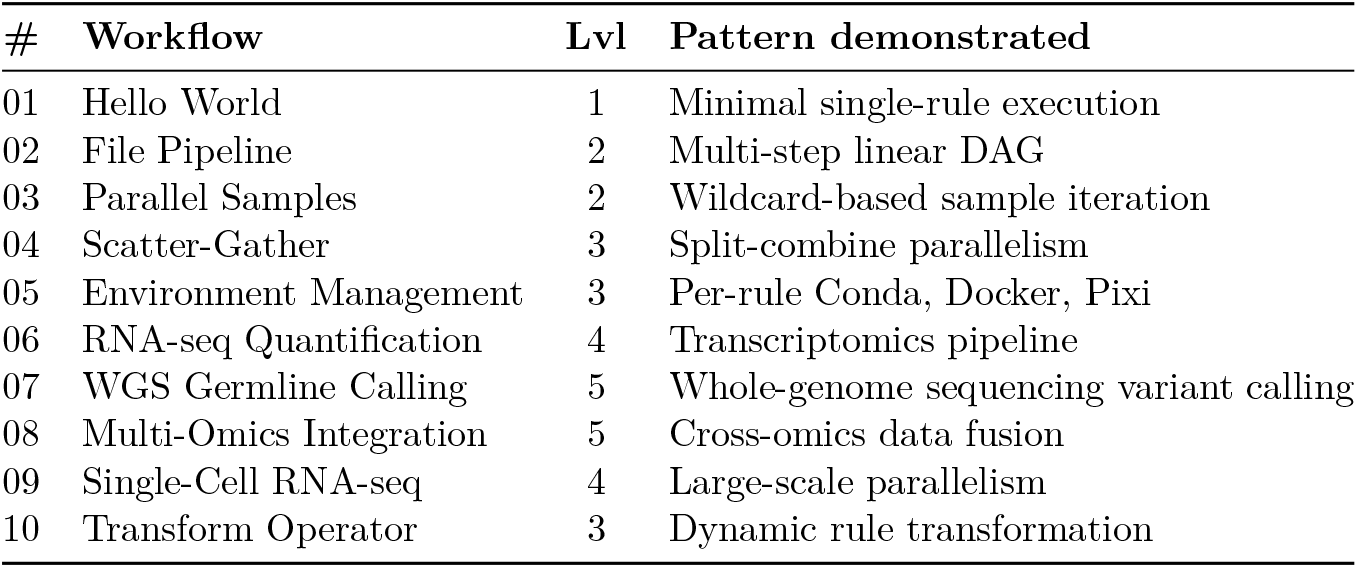
Workflow gallery. Complexity levels from 1 (single rule) to 5 (multi-omics integration). All examples pass validation and CI.

Three pipeline repositories exercise the full feature set in realistic contexts: `oxo-flow-circrna` (circular RNA detection), `oxo-flow-clindet` (somatic and germline variant detection across seven rule categories), and `oxo-flow-venus` (somatic variant calling with tumour-normal paired analysis). These pipelines use scatter-gather parallelism, environment groups, clinical reporting templates, and reference directory conventions. They are maintained as demonstration pipelines; independent clinical validation against reference truth sets has not been completed.

### 2.5 Reproducibility and comparison with existing systems

oxo-flow provides a layered reproducibility architecture (Figure 4). At the configuration layer, workflow specifications are normalised and hashed (SHA-256), producing a stable identifier independent of formatting changes. At the environment layer, each backend derives a content-addressed cache key from its specification (YAML content hash, Docker image digest, Pixi lockfile hash), guaranteeing that identical declarations produce identical runtime environments. At the execution layer, the DAG is constructed deterministically with stable topological ordering. At the verification layer, every output file is hashed and recorded in checkpoint state; the `provenance verify` command compares live checksums against stored records. The checkpoint JSON consolidates rule outcomes, benchmark records, and all checksums into a single-file audit trail that can be archived for regulatory compliance.

**Figure 4.**
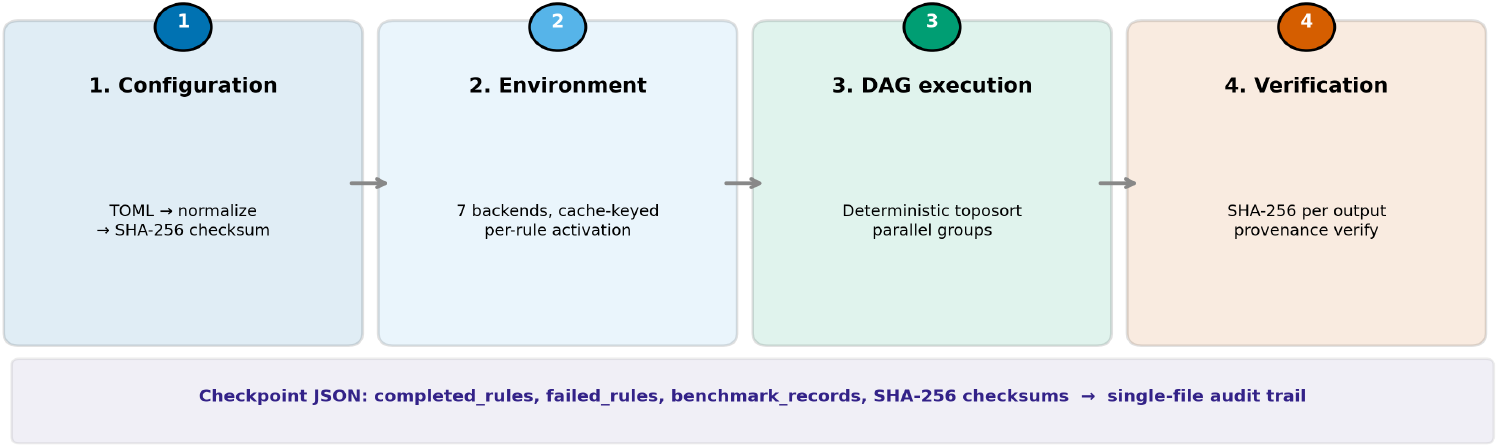
Layered reproducibility architecture.

Figure 5 compares capabilities across five workflow systems, verified against official documentation (June 2026). oxo-flow is the only system that combines a compiled native binary, memory-safety guarantees enforced at the compiler level, a declarative TOML format with JSON Schema validation, per-rule environment isolation across seven backends, a built-in web interface with REST API, clinical-grade reporting, command sanitisation, checkpoint-based fault tolerance with cryptographic output verification, and integrated container packaging. Dimensions where established systems lead—ecosystem size and community workflow collections, conditional workflow logic, and native cloud execution—are acknowledged.

**Figure 5.**
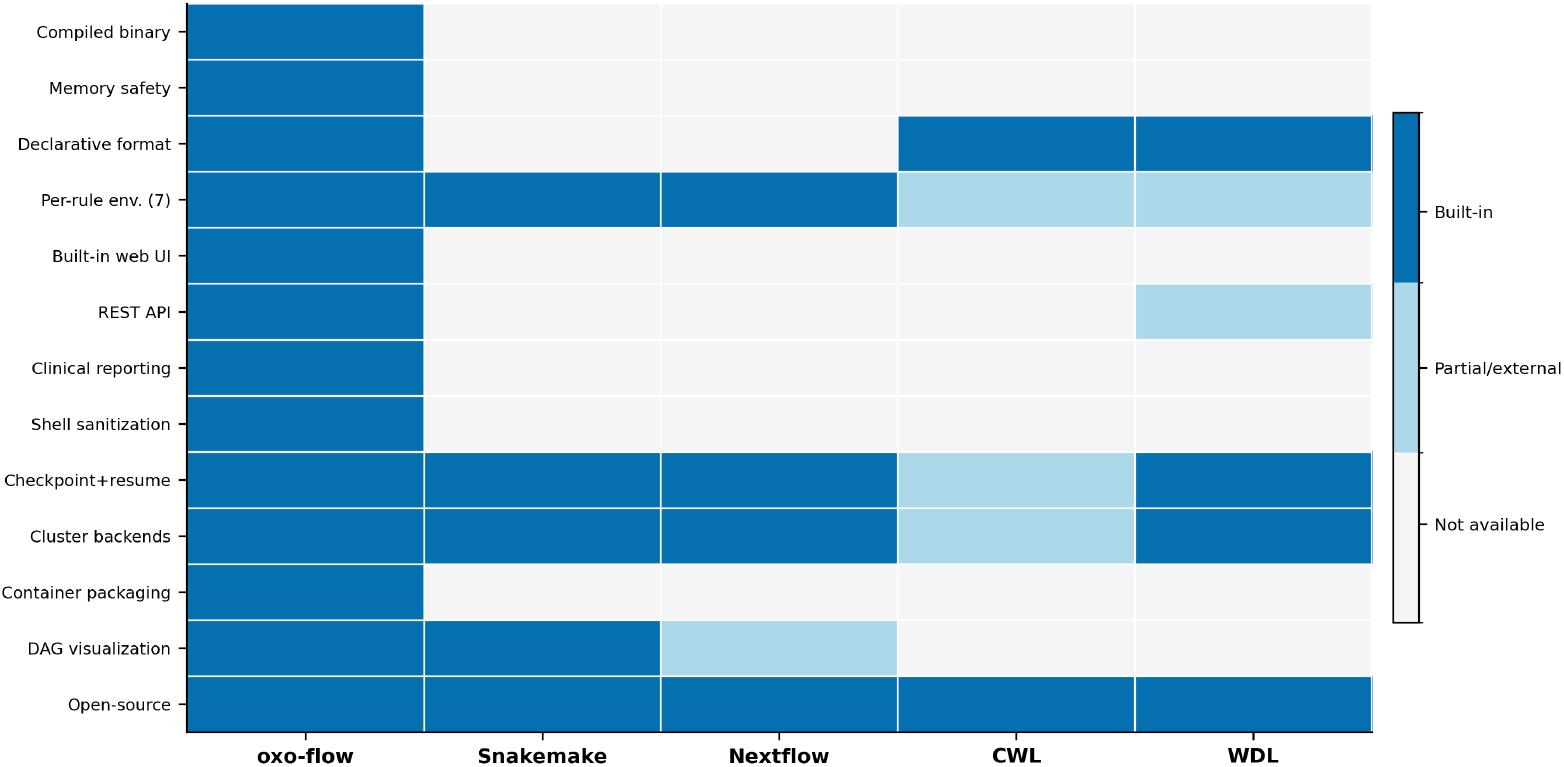
Capability matrix verified against official documentation (June 2026).

## 3 Discussion

oxo-flow occupies a previously empty position on the workflow engine design axis. Snakemake provides scripting flexibility through its Python DSL; Nextflow provides cloud-native scalability through reactive dataflow; CWL and WDL provide multi-engine portability. oxo-flow provides compile-time correctness for the engine itself—memory safety, type safety, and data-race freedom are guaranteed before the binary is ever executed. Workflow configurations are validated at parse time against a JSON Schema, catching malformed specifications before rule execution begins. These guarantees cover the engine and its configuration layer, not the bioinformatics tools being orchestrated or the biological validity of the analyses they perform.

The cost of this guarantee is expressiveness. oxo-flow workflows are declarative TOML, not executable scripts. A researcher cannot embed a Python function or Groovy closure in a workflow definition. For exploratory analyses, Snakemake or Nextflow may be more appropriate. For production genomics where reproducibility and auditability carry regulatory or clinical weight, the trade is warranted.

The latency results carry distinct implications for development versus production. During workflow development, a 13 ms validate-and-dry-run cycle enables an interactive feedback loop unavailable with Snakemake or Nextflow. The scaling data provide further assurance: workflow complexity imposes negligible additional planning overhead. For production runs measured in hours, the latency difference is immaterial; the memory efficiency becomes the more relevant advantage, enabling higher-density concurrent workflow validation on shared infrastructure.

Several limitations are genuine. oxo-flow is a young project—version 0.7.0, 401 commits, 19 releases—and its community cannot match the decade-plus maturity of Snakemake or Nextflow. It does not import existing CWL, WDL, or Snakemake workflows. The web interface provides a code editor rather than a visual DAG builder. A Kubernetes operator, native cloud object storage streaming, and Global Alliance for Genomics and Health (GA4GH) standards compliance are not yet implemented. The pipeline repositories are demonstration pipelines without independent clinical validation. Shell sanitisation, while providing meaningful protection, has documented limitations against encoded payloads and indirect injection vectors. As a young project, oxo-flow would benefit substantially from real-world testing and community feedback across diverse computing environments—cluster configurations, bioinformatics tool chains, and workflow patterns that cannot be fully replicated in continuous integration. The project actively encourages user reports of both successes and failures to guide development priorities.

## 4 Methods

### 4.1 Implementation

oxo-flow is implemented in Rust (2024 edition) with the following design properties: all crates enforce `#![forbid(unsafe_code)]`, guaranteeing the absence of memory-unsafe code; the `WorkflowConfig` type uses a type-state pattern (Parsed *→* Ready *→* Validated) to prevent invalid state transitions at compile time; and wildcard expansion uses a lazily-compiled regular expression to avoid repeated compilation. Core dependencies: petgraph 0.8 [21] (DAG operations via `toposort`), Tokio 1 [23] (asynchronous runtime), Clap 4.5 (command-line interface), Axum 0.8 [22] (web server), Serde 1 (serialisation), Tera 1 (HTML templating), Reqwest 0.12 (HTTP client), SQLx 0.8 [24] (database), sysinfo 0.38 (resource detection). Release builds use optimisation level 3, full link-time optimisation (LTO), single codegen unit, and symbol stripping. CI enforces formatting, linting with zero-tolerance for warnings, compilation, and test execution on every commit.

Shell command safety is provided by four validation functions that inspect every command before execution: removal of command chaining operators and subshell expansion constructs, detection of injection patterns, prevention of wildcard-mediated injection, and blocking of path traversal attempts. Interpreter paths for 14 script extensions are validated against a configurable allowlist. Documented limitations of this approach include: encoded payloads (Base64, `python -c`, `perl -e`) are not detected; URL-encoded path traversal sequences bypass detection; wildcard values are substituted after sanitization. The project recommends using regex-based wildcard constraints to restrict allowed characters in user-supplied values.

### 4.2 Benchmarks

All benchmarks are produced by a single reproducible script recording hardware, software versions, and raw measurement data. Latency benchmarks used hyperfine 1.20.0 [25] with documented warmup and run counts. Wildcard benchmarks used Criterion.rs (3 s warmup, 100 samples, outlier detection). DAG benchmarks used linear chain graphs of 10–1,000 nodes. Memory measurements used `/usr/bin/time -l`. All measurements on Apple M5 MacBook Pro (10 cores, 24 GB unified memory, macOS 26.5.1). Snakemake 9.22.0 was installed via Conda; Nextflow 26.04.3 via the official installation script with OpenJDK 17.0.18. Cross-tool comparisons measured Snakemake `--help` and Nextflow `-version` (binary startup) against oxo-flow dry-run (full workflow parse-to-plan cycle); matched dry-run benchmarks across DSLs were not performed.

### 4.3 Data availability

Source code, documentation, workflow examples, and test data are available at https://github.com/Traitome/oxo-flow under Apache License 2.0 (core and CLI) and dual academic/commercial licensing (web server). oxo-flow can be installed via Cargo (`cargo install oxo-flow-cli`) or via Bioconda (`conda install -c bioconda oxo-flow-cli`); pre-built binaries are available from GitHub Releases. Pipeline repositories: https://github.com/Traitome/oxo-flow-circrna, https://github.com/Traitome/oxo-flow-clindet, https://github.com/Traitome/oxo-flow-venus. Benchmark scripts and raw measurement data are included as supplementary material.

### 4.4 Author contributions

S.W. conceived the project, designed the architecture, implemented the engine, CLI, and web server, conducted all benchmarks, and wrote the manuscript.

### 4.6 Funding

Central South University Startup Funding; National Natural Science Foundation of China (82303953); Hunan Provincial Natural Science Foundation of China (2025JJ40079).

## References

[1] J. Köster and S. Rahmann, “Snakemake—a scalable bioinformatics workflow engine,” Bioinformatics, vol. 28, no. 19, pp. 2520–2522, 2012.

[2] P. Di Tommaso, M. Chatzou, E. W. Floden, P. P. Barja, E. Palumbo, and C. Notredame, “Nextflow enables reproducible computational workflows,” Nat. Biotechnol., vol. 35, no. 4, pp. 316–319, 2017.

[3] B. Grüning, R. Dale, A. Sjödin, et al., “Bioconda: sustainable and comprehensive software distribution for the life sciences,” Nat. Methods, vol. 15, no. 7, pp. 475–476, 2018.

[4] F. Strozzi, R. Janssen, R. Wurmus, et al., “Scalable workflows and reproducible data analysis for genomics,” in Evolutionary Genomics, Methods Mol. Biol., vol. 1910, pp. 723–745, Springer, 2019.

[5] F. Mölder, K. P. Jablonski, B. Letcher, et al., “Sustainable data analysis with Snakemake,” F1000Research, vol. 10, p. 33, 2021.

[6] S. Kieser, J. Brown, E. M. Zdobnov, M. Trajkovski, and L. A. McCue, “ATLAS: a Snake-make workflow for assembly, annotation, and genomic binning of metagenome sequence data,” BMC Bioinform., vol. 21, no. 1, p. 257, 2020.

[7] L. Czech and M. Exposito-Alonso, “grenepipe: a flexible, scalable and reproducible pipeline to automate variant calling from sequence reads,” Bioinformatics, vol. 38, no. 20, pp. 4809–4811, 2022.

[8] B. Chapman, J. Chilton, M. Heuer, et al., “Common Workflow Language (CWL), v1.0,” Figshare, 2016.

[9] K. Voss, G. Van der Auwera, and J. Gentry, “Full-stack genomics pipelining with GATK4 + WDL + Cromwell,” F1000Research, vol. 6, no. ISCB Comm J, p. 1381, 2017.

[10] K. A. Kyritsis, N. Pechlivanis, and F. Psomopoulos, “Software pipelines for RNA-Seq, ChIP-Seq and germline variant calling analyses in Common Workflow Language (CWL),” Front. Bioinform., vol. 3, p. 1275593, 2023.

[11] P. K. Korhonen, R. S. Hall, N. D. Young, and R. B. Gasser, “Common Workflow Language (CWL)-based software pipeline for de novo genome assembly from long-and short-read data,” GigaScience, vol. 8, no. 4, p. giz014, 2019.

[12] F. Z. Khan, S. Soiland-Reyes, R. Sinnott, et al., “CWLProv: Interoperable retrospective provenance capture and computational analysis sharing,” in Proc. WORKS 2019, IEEE, 2019.

[13] E. Afgan, D. Baker, B. Batut, et al., “The Galaxy platform for accessible, reproducible and collaborative biomedical analyses: 2018 update,” Nucleic Acids Res., vol. 46, no. W1, pp. W537–W544, 2018.

[14] M. Tekman, B. Batut, A. Ostrovsky, et al., “A single-cell RNA-sequencing training and analysis suite using the Galaxy framework,” GigaScience, vol. 9, no. 10,.giaa102, 2020.

[15] E. C. Anderson, S. R. Piccolo, Z. E. Ence, et al., “Simplifying the development of portable, scalable, and reproducible workflows,” eLife, vol. 10, p. e71069, 2021.

[16] J. Ding, Y. Peng, R. Wei, B. Wang, J.-G. Zhou, and S. Wang, “BLIT: an R package for seamless integration of command-line bioinformatics tool universe,” Bioinform. Adv., vol. 6, no. 1, p. vbag088, 2026.

[17] B. Chen, D. Wu, and G. Zhang, “GFFx: A Rust-based suite of utilities for ultra-fast genomic feature extraction,” GigaScience, vol. 14, p. giaf124, 2025.

[18] S. Vijendran, T. K. Anderson, A. Markin, and O. Eulenstein, “Phylo-rs: an extensible phylogenetic analysis library in rust,” BMC Bioinform., vol. 26, p. 197, 2025.

[19] N. D. Matsakis and F. S. Klock II, “The Rust language,” in Proc. HILT 2014, ACM, pp. 103–104, 2014.

[20] T. Preston-Werner, “TOML v1.0.0,” y2021. https://toml.io

[21] petgraph v0.8. https://crates.io/crates/petgraph

[22] axum v0.8. https://github.com/tokio-rs/axum

[23] tokio v1. https://tokio.rs

[24] sqlx v0.8. https://github.com/launchbadge/sqlx

[25] hyperfine v1.20.0. https://github.com/sharkdp/hyperfine

[26] Traitome, “oxo-flow: A Rust-native bioinformatics pipeline engine,” v0.7.0, 2026. https://github.com/Traitome/oxo-flow

