## supplemental data info for "oxo-flow: compiled, memory-safe bioinformatics workflow orchestration": README.md

# Raw Measurement Data — oxo-flow Paper

All measurements taken 2026-06-10/11. Every number in the manuscript traces to
a file in this directory or to the oxo-flow source tree.

## Hardware Under Test

```
Model:       Apple M5 MacBook Pro (Mac17,2)
Processor:   Apple M5, 10 cores (4 performance + 6 efficiency)
Memory:      24 GB unified memory
OS:          macOS 26.5.1 (Build 25F80)
Arch:        arm64
```

## Measurement 1: Startup Latency

**File**: `startup_hyperfine.json`
**Tool**: hyperfine 1.20.0
**Command**: `hyperfine --warmup 10 --runs 50 "oxo-flow --help > /dev/null"`
**Result**: mean 5.94 ms, σ 0.28 ms, median 5.92 ms, range 5.36–6.79 ms

**Comparison — Snakemake**:
**Version**: Snakemake 9.22.0 (conda environment bench_snakemake)
**Command**: `hyperfine --warmup 3 --runs 20 "snakemake --help > /dev/null"`
**Result**: mean 647.4 ms, σ 15.0 ms, range 624.5–684.8 ms
**Speedup**: 647.4 / 5.94 = 109×

## Measurement 2: Wildcard Benchmarks

**File**: `criterion_wildcard_raw.txt`
**Tool**: Criterion.rs (`cargo bench -p oxo-flow-core`)
**Method**: 3 s warmup, 100 samples, outlier detection
**Bench 1 — expand_pattern**: 591.80–594.53 ns (mean 593.12 ns)
  Pattern: `results/{sample}/{chr}/reads_R{read}.fastq.gz`
**Bench 2 — cartesian_expand_large**: 24.649–24.819 µs (mean 24.734 µs)
  Combinations: 10 samples × 2 reads × 22 chromosomes = 440

## Measurement 3: Binary Sizes

**CLI** (`target/release/oxo-flow`): 9.3 MB stripped
**Web** (`target/release/oxo-flow-web`): 5.9 MB
**Installed** (`~/.cargo/bin/oxo-flow`): 16 MB unstripped

## Measurement 4: Code Metrics

**Core library**: 24,417 lines across 25 source files
**CLI crate**: 4,618 lines across 14 source files  
**Web crate**: 6,554 lines across 20 source files
**Total Rust files** (all crates): 61
**Test annotations** (core): 583 #[test] + 15 #[tokio::test] = 598

## Measurement 5: Version History

**Commits**: 401 (`git rev-list --count HEAD`)
**Releases**: 19 (`git tag | wc -l`)
**Latest**: v0.7.0
**Earliest**: v0.1.0

## Measurement 6: CLI Commands

**Count**: 31 subcommands
**Source**: `crates/oxo-flow-cli/src/main.rs` (Clap derive enum)

## Measurement 7: REST API Endpoints

**Count**: 60 routes
**Source**: `crates/oxo-flow-web/src/lib.rs` (axum Router)
**Handlers**: 13 handler modules under `handlers/`

## Measurement 8: DAG Construction

**Method**: Linear chain stress test (dag.rs)
**Results**:
  - 10 nodes: 0.18 ms
  - 50 nodes: 0.42 ms
  - 100 nodes: 0.85 ms
  - 500 nodes: 5.2 ms
  - 1,000 nodes: 12.8 ms

## Measurement 9: Security Functions

**Source**: `crates/oxo-flow-core/src/executor/security.rs` (302 lines)
**Functions**: sanitize_shell_command (L159), validate_shell_safety (L197),
  validate_wildcard_injection (L225), validate_path_safety (L262)
**Verified**: imported and called from executor/process.rs and executor/hooks.rs

## Measurement 10: Cross-Compilation

**File**: `Cross.toml`
**Targets**: aarch64-unknown-linux-musl, armv7-unknown-linux-musleabihf,
  x86_64-unknown-linux-musl
**CI targets**: x86_64-apple-darwin, aarch64-apple-darwin
